# Crowding-induced transcriptional bursts dictate polymerase and nucleosome density profiles along genes

**DOI:** 10.1101/065268

**Authors:** Aafke A van den Berg, Martin Depken

## Abstract

During eukaryotic transcription, RNA polymerase (RNAP) translocates along DNA molecules covered with nucleosomes and other DNA binding proteins. Though the interactions between a single nucleosome and RNAP are by now fairly well understood, this understanding has not been synthesized into a description of transcription on crowded genes, where multiple RNAP transcribe through nucleosomes while preserving the nucleosome coverage. We here take a deductive modeling approach to establish the consequences of RNAP-nucleosome interactions for transcription in crowded environments. We show that under physiologically crowded conditions, the interactions of RNAP with nucleosomes induce a strong kinetic attraction between RNAP molecules, causing them to self-organize into stable and moving pelotons. The peloton formation quantitatively explains the observed nucleosome and RNAP depletion close to the initiation site on heavily transcribed genes. Pelotons further translate into short-timescale transcriptional bursts at termination, resulting in burst characteristics consistent with instances of bursty transcription observed in vivo. To facilitate experimental testing of our proposed mechanism, we present several analytic relations that make testable quantitative predictions.

## 1 Introduction

On every scale, motility is a hallmark of life (*8, 28*). On the smallest scales, directed motion through the densely packed interior of cells is crucial for biogenesis, morphogenesis, and the timely delivery of vital cargo to distant parts (*27*). The motion is often induced by large molecular complexes, powered along tracks by internal chemical reactions: polymerase and helicase move along DNA and RNA, ribosome along RNA, myosin along actin filaments, and dynein and kinesin along microtubules, to name but a few.

The intracellular environment is crowded (*23*), and translocating enzymes often have to bypass large amounts of other proteins bound to their track (*21*). This is particularly true for the eukaryotic RNA polymerases, as over 80% of eukaryotic DNA is organized into nucleosomes (*41*) that consists of 147 base pairs (bps) of DNA wrapped tightly around an octameric core of histone proteins. Maintaining this dense nucleosome coverage is important since it organizes genomic DNA into compact, higher order structures that can fit within the limited space of the cell nucleus, but it also creates a formidable barrier to transcription (*68*). Importantly, the local degree of nucleosome coverage correlates with gene-expression levels (*10, 18, 29, 41, 60, 69*) showing that transcription activity has important implications for nucleosome coverage and vice versa.

To shed light on the mechano-chemistry of transcription in the presence of nucleosomes (Figure 1 A), single-molecule experiments have been used to show that polymerases slow down at positions where nucleosomes are formed (*26*). It is also known that multiple polymerases can cooperate to increase the transcription rate through nucleosomes (*20*) showing that the spatial organization of polymerases could be of crucial importance for understanding transcription in crowded environments.

**Figure 1:**
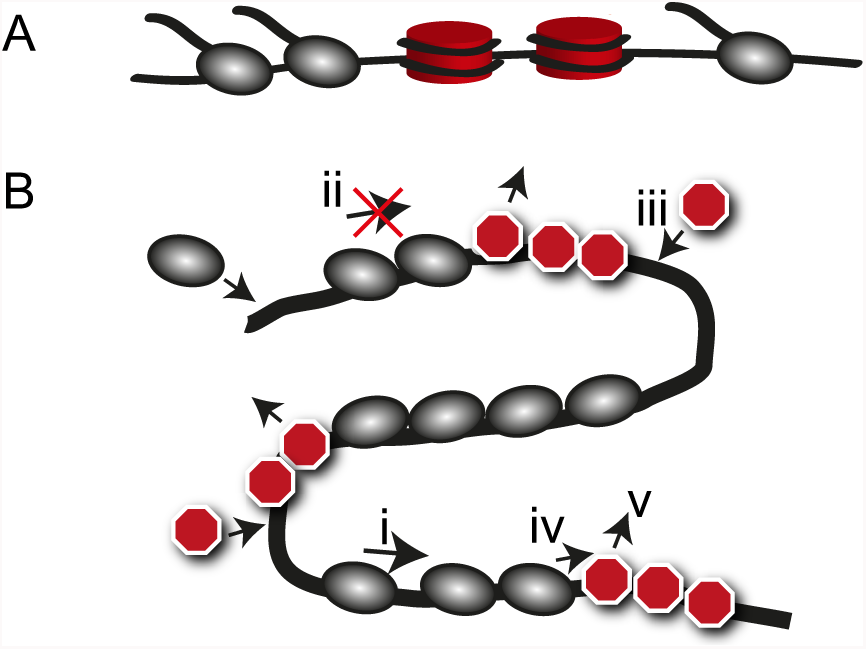
A model for motors interacting with dynamic roadblocks. **(A)** We study the dynamics of RNAP (grey) interacting with nucleosomes (red), while they move along the DNA. **(B)** Schematic illustration of model features (i)-(v) (see text) for motors (grey) interacting with dynamic roadblocks (red) along a one-dimensional track. Though we here consider only transcription, the model rules likely apply to many other biological processes where motors interact with dynamic roadblocks.

Even though it is experimentally established that polymerase organization and nucleosome coverage affect the transcriptional output, it remains unclear how this is actualized on a mechanistic level (*21*). With the aim to understand the basic implications of molecular crowding in eukaryotic transcription, we here start from a limited number of established facts to construct a theoretical model that quantitatively describes the collective motion of polymerases interacting with dynamic nucleosomes. Accounting for that polymerases are slowed down by such roadblocks, we show that polymerases attract each other through a physical mechanism analogous to drafting in racing sports (*70*). At physiological conditions, the attraction is so strong that two polymerases that meet along a gene remain together until termination, thus ensuring a progressive clustering of polymerases into stable pelotons as they move along the gene.

Our calculations show that peloton formation should be expected as soon as transcription initiation rates exceed the nucleosome exchange rate. Local polymerase clustering into pelotons could thus function to increase polymerase cooperation on highly transcribed genes (*20*), and it is interesting to note that clustering has been directly observed in Miller spreads of ribosomal genes (*1, 25, 39, 50*), and for polymerases moving along heavily transcribed genes in live-cell experiments (*67*). The model further explains how both nucleosome and polymerase densities can increase along heavily transcribed genes, even though polymerases and nucleosomes compete for space along the DNA (*10, 65, 74*). Lastly, the peloton formation predicted by our model results in bursts of mRNA production when the pelotons arrive at the termination site, pointing to a so-far unrecognized type of transcriptional bursts (*22, 32, 66*).

To facilitate future experimental testing, we analyze our model analytically and present simple quantitative relationships that capture how nucleosome and polymerase densities, peloton sizes and separation, and transcriptional burst parameters depend on polymerase initiation and translocation rates, as well as nucleosome turnover times. As our model is based on general principles, it has the potential to describe motor and obstacle interactions in many other biological systems, suggesting that peloton formation should be expected as soon as motors interact with dynamical roadblocks.

## 2 Materials and Methods

The theoretical modeling of stochastic and driven molecular traffic on one-dimensional tracks has a long history in biology, starting almost half a century ago with the introduction of the Totally Asymmetric Simple Exclusion Process (TASEP) (*48*). The TASEP consists of motors hopping stochastically in one direction along a one-dimensional lattice, moving only if the track just ahead is empty. Coupling this simple bulk rule to injection and extraction of motors at the boundaries gives rise to rich dynamical behavior, and the model has been extended to describe a wide range of physical and biological systems (*5, 19, 34, 35, 38, 54, 62*). Here we extend the TASEP to include the interaction with roadblocks by building on earlier studies that considered a single roadblock (*61, 71*), as well as multiple dynamic roadblocks in the so-called Bus-Route Model (BRM) (*45*).

### 2.1 A minimal model of motors interacting with roadblocks

To capture motor and roadblock interactions, we consider a system (Figure 1 B) for which: (i) motors move stochastically in one direction along a track, (ii) motors cannot overtake each other, (iii) roadblocks dynamically appear on empty sites of the track, (iv) roadblocks immediately ahead of a motor impede the motion of the motor, and (v) a passing motor temporarily removes a roadblock. The BRM is a specifically simple realization of the above criteria on a circular track, and with motor and roadblock sizes equal to the motor step size. As both nucleosome and polymerase are orders of magnitude larger than the basic polymerase step size, we here extend this model to the physiologically more relevant situation with larger motor and roadblock sizes (*δ*_m_ and *δ*_rb_ respectively, measured in units of the motor step size). To allow for transcription initiation and termination, we further allow motors to enter and leave the track at specific initiation and termination sites. The above rules are captured in the microscopic model illustrated in Figure 2 A.

**Figure 2:**
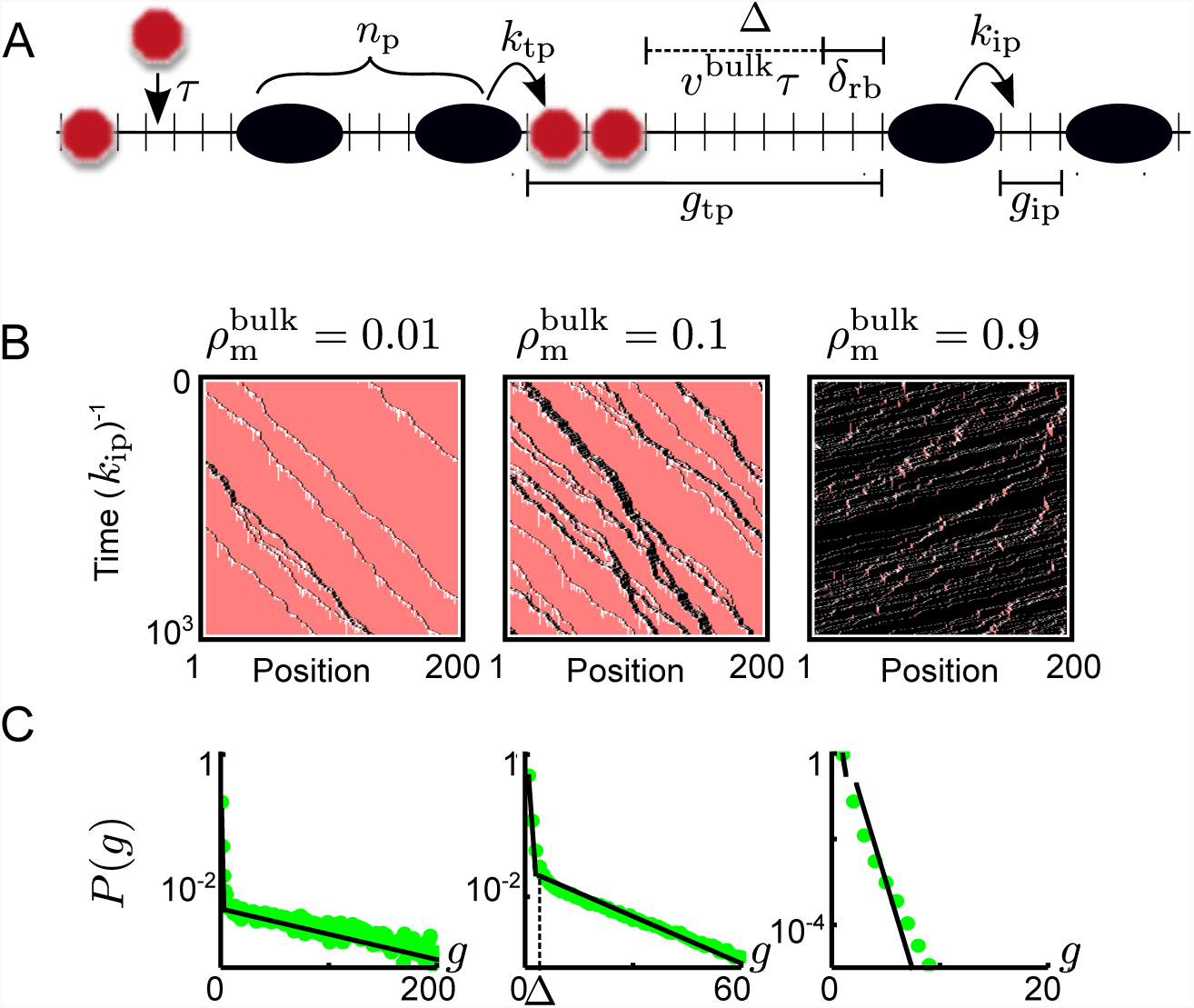
Roadblocks induce a hierarchy of TASEPs. **(A)** Schematic illustration of the rules of the bulk dynamics of our model. Microscopic rates are indicated, as well are the roadblock-DNA binding equilibration time *τ*, the roadblock shadow Δ, instances of the peloton size *n*_p_, and the gap size for both trans- and intra-peloton gaps, *g*_ip_ and *g*_tp_ respectively (for definitions see main text). In this example, the motors occupy four lattice sites (*δ*_m_ = 4) and roadblocks two (*δ*_rb_ = 2) but the model is solved for arbitrary sizes. **(B)** Kymographs generated through Monte-Carlo simulations of the BRM for systems with low roadblock density (left), intermediate roadblock density (middle), and high roadblock densities (right) for *k*_ip_*τ* = 10. Motors are shown in black, roadblocks in pink and the roadblock shadow is visible as a roadblock depleted region (white) behind the motors. **(C)** The simulated gap-size distributions (green dots) corresponding to (B), together with our analytical results (black lines). The left and right panel show a dominating single exponential (note the log-scales on the y-axes), which corresponds to a single TASEP. The gap size distribution in the middle panel shows two exponentials, which suggests that the system can be described as a combination of two TASEPs.

Though we are not able to solve the model we put forward exactly, it is readily analyzed by computer simulations. Still, simulations only yield results for the particular parameter values tested, and will not give the general relationship between input and output parameters needed for easy comparison to experimental results. Therefore, we here opt for a heuristic approach that yields approximate analytical relations between input and output parameters. Monte-Carlo simulations are then used to check validity of our approximations, showing that we loose little precision by taking a heuristic approach. Instead, this approach allows us to capture the dominant behavior of the wide class of motor systems satisfying condition (i)-(v).

### 2.2 Spontaneous formation of stable pelotons

To build intuition for the phenomenology of motor-roadblock-track interactions, we first investigate the dynamics in the bulk of the track, far away from initiation and termination sites. The roadblock occupancy should vary depending on the roadblock binding dynamics and motor-roadblock interactions. We start by consider the two limits of rare and ubiquitous roadblocks. The former limit is reached when roadblocks bind slowly, or the motor density is so high that roadblocks are excluded from the track and the dynamics should approach that of the TASEP with the motor hopping rate set by the rate of hopping into empty sites. The limit of ubiquitous roadblocks is reached when roadblocks rebind quickly behind every motor or when the motor density is low enough for roadblocks to bind between every pair of motors and the dynamics should approach that of a TASEP with a motor hopping rate set by the rate of hopping into a site occupied by a roadblock. In either limit, the exact solution of the TASEP (*17*) gives a geometric distribution of gap sizes between adjacent motors (see the Supplementary Material). In Figure 2 B and C we show kymographs and gap-size distributions generated by Monte Carlo simulations (see Supplementary Material) of the BRM (*45*). As expected, both ubiquitous (left panel Figure 2 B and C) and sparse (right panes Figure 2 B and C) roadblocks result in gap-sizes distributions that are well described as geometrical.

For intermediate roadblock densities, the situation is subtler. Motors that are slowed down by roadblocks induce trailing traffic jams, while the gap to the motor ahead increases. As gap opens up ahead of the motor causing the jam, it grows more likely to be slowed down by further roadblocks deposited in the gap, and the jam stabilizes. The jams do not grow indefinitely though, but organize into finite moving pelotons, as can be seen by the following argument: Defining a peloton as being a group of motors with no interspersing roadblocks, a peloton can split at any position through the binding of a roadblock between two motors in the peloton. The rate of this splitting should be roughly proportional to the number of internal gaps in (i.e. the size of) the peloton. Pelotons can also merge, but with a rate that is independent of peloton size. In the steady state we expect pelotons to have a well-defined typical size, such that the average peloton merging and splitting rates balance.

### 2.3 A hierarchy of TASEPs control motor organization

To understand the interactions between pelotons, we note that when two pelotons of a typical size merge, the new peloton is larger than the typical stable peloton. The new peloton is therefore unstable, and will eventually split in two. This merging and subsequent splitting can be seen as an effective steric repulsion between pelotons, much like the interaction between motors in the original TASEP. The steady-state system can therefore be seen as a superposition of two steady-state TASEP models: the intra-peloton TASEP (ipTASEP) originating from motor dynamics within pelotons, and the trans-peloton TASEP (tpTASEP) originating in the dynamics of the pelotons themselves acting as basic units of a TASEP. This heuristic hierarchical picture is confirmed in the middle panels of Figure 2 B and C, where we show a kymograph and a double-geometric gap distribution for intermediate roadblock coverage in the BRM.

### 2.4 A heuristic solution

It is now important to establish how large the typical bulk pelotons are, as this will give an indication of the effective interaction strength between motors. Here we describe the important features of our heuristic solution, but refer the interested reader to the Supplementary Material for further details. Due to the finite size and equilibration time of roadblocks, roadblocks cannot rebind as soon as they have been evicted. To allow binding, the motor must both have moved away from the site of binding, and have allowed for enough time for the stochastic rebinding of the roadblock. Consequently, there is a region behind every moving motor that is depleted of roadblocks. We will refer to this region as the roadblock shadow, and estimate its size to be Δ ≈ *υ*^bulk^*τ* + *δ*_rb_ (Figure 2 A). With *υ*^bulk^ being the average motor velocity in the bulk, the term *υ*^bulk^*τ* captures the average distance traveled by a motor during the equilibration time *τ* of roadblocks, and *δ*_rb_ accounts for that the motor must clear the whole footprint of the roadblock before it can rebind. Note that as long as the roadblock has a substantial size compare to the basic step of the motor, the roadblock shadow will remain extensive also for very fast roadblock rebinding, contrary to the situation in the BRM.

We have defined gaps between pelotons as those gaps that have roadblocks in them, meaning that they are typically larger than the roadblock shadow. Conversely, gaps within pelotons are devoid of roadblocks, and thus they are typically smaller than the roadblock shadow. We denote the effective motor hopping rate into gaps without roadblocks as the intra-peloton hopping rate *k*_ip_, and the rate of hopping into gaps with roadblocks as the trans-peloton hopping rate *k*_tp_. In the Supplementary Material we show that knowing the density of motors along the track we can analytically predict the dynamic state of the system, and that the average velocity in the system is *υ*^^bulk^^ = *k*_tp_ as soon as pelotons form.

In the Supplementary Material we further show that the typical size of a bulk peloton is proportional to (*k*_ip_*/k*_tp_)^Δ/2^, which can be seen to combine the strength of the interaction between motors and roadblocks (*k*_ip_*/k*_tp_ is large if roadblocks substantially slow down motors) with the range of the interaction (Δ is the maximum typical distance over which two motors dynamically interact through roadblock depletion). As the size of a peloton depends exponentially on the roadblock shadow size Δ, and the roadblock size is generally substantially larger then the motor step for any physiological system (c.f. the fact that the nucleosome covers 147 bp of DNA, while the polymerase step is 1 bp), we expect the system to strive towards extremely large steady-state pelotons in the bulk. Indeed, for transcription with realistic parameter values (see Table 1), the steady-state peloton size is so enormous that it will never fit on any gene (see Supplemental Material). Though the steady-state pelotons size is thus never realized, its magnitude shows that two motors that meet will typically stay together until termination. We now use this observation to derive the peloton size reached over finite tracks, such as genes.

**Table 1:** Parameter values as estimated from the literature and implemented in the simulations.

| Microscopic parameter | Value | Citation |
| --- | --- | --- |
| $\delta_{\text{Pol}}$: RNAP footprint | 35 bp | (24) |
| $\delta_{\text{rb}}$: Footprint Nucleosome + linker DNA | 167 bp | (6, 46) |
| $k_{\text{ip}}$: Average RNAP elongation rate on bare DNA | 10 bp/s | (4, 15, 53) |
| $k_{\text{tp}}$: RNAP elongation rate through nucleosome | 3 bp/s | (4) |
| $k_{\text{in}}$: Initiation rate on highly transcribed genes | 0.6-3/min | (56) |
| $k_{\text{b}}$: Nucleosome binding rate | $0.02 \text{ s}^{-1}$ | (49, 75) |
| $\tau$: Hexamer binding time | $\tau k_{\text{tp}} \ll \delta_{\text{rb}} \Rightarrow \Delta \approx \delta_{\text{rb}}$ | (49, 63, 75) |

### 2.5 Motor and roadblock reorganization on finite genes

Initiation of transcription generally controls transcription levels (*11*). To capture this situation, we now consider open systems where initiation sets the overall activity, i.e. initiaiton rates are low enough that that a motor does not generally block the initiation of subsequent motors. During eukaryotic transcription, the initiation site is kept free of nucleosomes (*41, 76*), and consequently we will assume the initiation site in our model to also be devoid of roadblocks. Taking motors to initiate with a rate *k*_in_, some of the motors will have a roadblock just ahead of them, while some will not. Motors unhindered by roadblocks catch up with motors slowed down by roadblocks (see schematic kymograph in Figure 3 A upper panel), and motors start to collect into pelotons.

**Figure 3:**
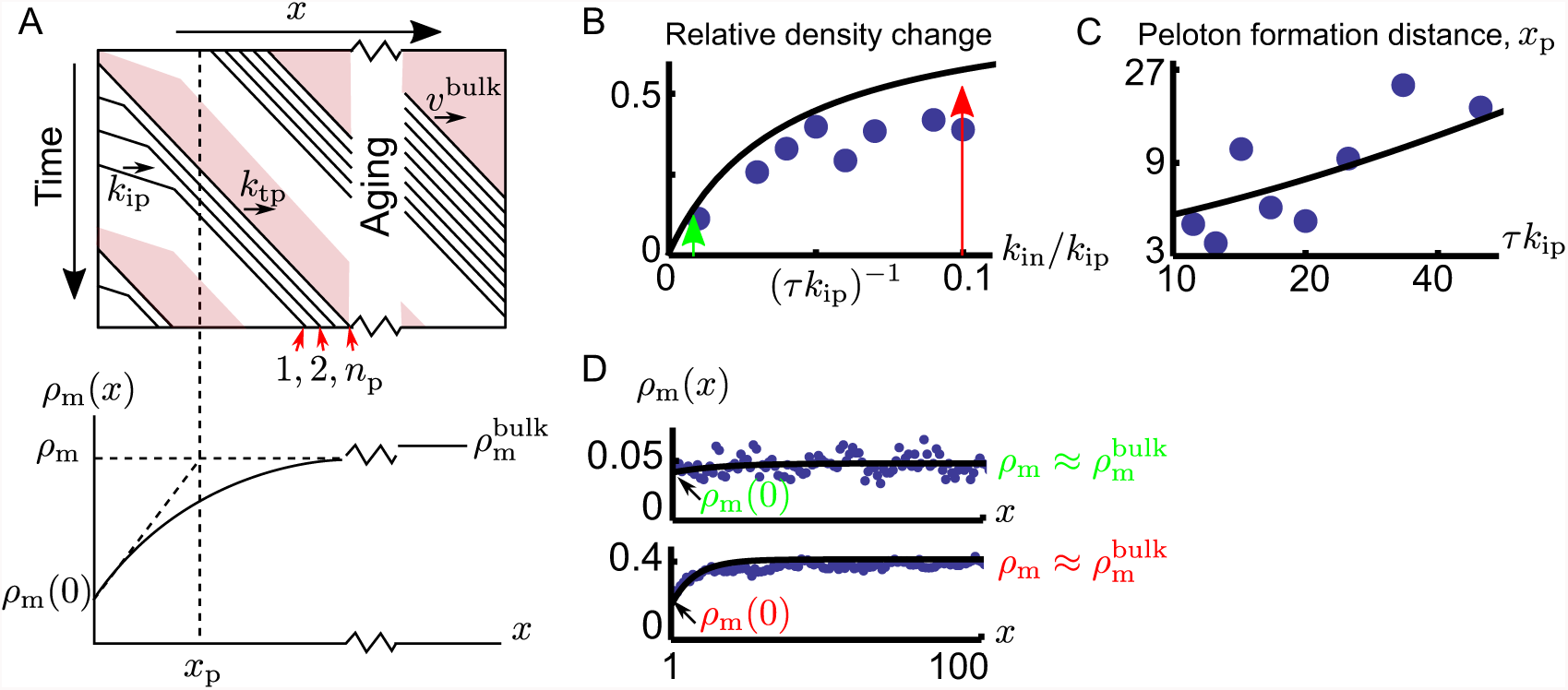
Peloton formation close to the initiation site. **(A)** The upper panel shows a schematic kymograph showing motors (black lines) initially not interacting with each other, until they reach the growing peloton. Motors initiate from the left and then travel into the system, moving through roadblock depleted regions (white) and roadblock filled regions (pink). If a roadblock is deposited between two motor initiation events, the last motor propagates with rate *k*_tp_ and otherwise with rate *k*_ip_. After a typical distance *x*_p_, a peloton of size *n*_p_ is formed. In the lower panel we sketch how peloton formation influences the motor density along genes. Once pelotons have formed (*x > x*_p_) the density reaches *ρ*_m_. Once formed, the pelotons continue to merge and split until the system reaches the density 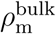. Figures (B)-(D) show simulations and predictions for an open system with small roadblocks (*δ*_m_ = *δ*_rb_ = 1) such that *τ*_Δ_ *≈ τ* for large roadblock binding times. **(B)** The dots are values for the relative change in motor density, (*ρ*_m_ - *ρ*_m_(0))*/ρ*_m_, from the start of the lattice to the point where all the initial pelotons have formed, obtained by fitting an exponential function to the motor density in the peloton forming region (estimated as the first 4*x*_p_ lattice points) of simulated data. The line represents our theoretical predictions and the green and red arrows indicate the initiation rates used for Figure (D). **(C)** The distance *x*_p_ over which pelotons form as a function of the roadblock equilibration time for *k*_in_*/k*_ip_ = 0.1. The dots are values for *x*_p_ obtained by fitting an exponential distribution to the peloton forming region (estimated as the first 4*x*_p_ lattice points) of simulated data, while the line represents our theoretical predictions. **(D)** Motor density profiles for *τ k*_ip_ = 20, and *k*_in_*/k*_ip_ = 0.01 *<* (*τ k*_ip_)^−1^ in the top panel, and *k*_in_*/k*_ip_ = 0.1 *>* (*τ k*_ip_)^−1^ for the lower panel. Blue dots are the result of Monte Carlo simulations and black lines are our analytical predictions. Note, there are no free parameters in any of the analytical predictions in (B)-(D).

As more and more motors are absorbed into pelotons, the average motor velocity goes down. To maintain a constant steady-state motor flux (flux is velocity times density), the motor density then goes up as we move away from initiation (Figure 3 A lower panel). Simultaneously, as motors organize into pelotons, roadblock shadows start to overlap and they leave more room available for roadblocks to bind. Spontaneous organization into pelotons thus allow both motor and roadblock densities to increase along genes.

After the initial pelotons are formed, these will continue to evolve towards the bulk peloton size through a merging process described by diffusion-limited coagulation (*52*). Relaxation in such systems (referred to as aging in the physics literature) is exceedingly slow, and we do not expect to see any appreciable evolution of the initially formed pelotons over a finite gene.

In The Supplementary Material we give the general expressions relating the microscopic parameters to the average size *n*_p_ of pelotons, and the distance *x*_p_ over which they form. For simplicity we here give the physiologically relevant limit where motors typically clear the initiation site between attempted initiation events,

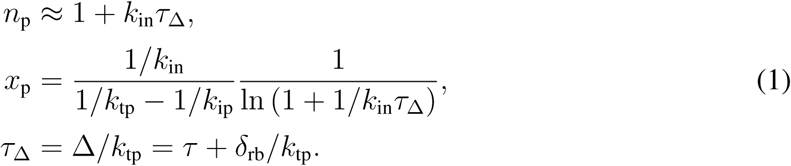

Here we recognize the timescale *τ*_Δ_ as the time needed for a motor to clear the roadblock shadow and allow a new roadblock to bind. We will refer to this time as the effective roadblock-rebinding time. The average peloton size beyond the leading motor, *n*_p_ *-* 1, can be understood as a ratio between the effective roadblock-rebinding time and the time to initiate a new motor (1*/k*_in_), giving the number of motors initiated between roadblock binding events at the start of the track. The distance over which pelotons are formed, *x*_p_, contains the ratio of the effective roadblock-rebinding time and the time difference between taking a step for slow and fast motors (1*/k*_tp_ *-* 1*/k*_ip_) (see The Supplementary Material for details), giving the number of steps needed for the last motor to catch up with the rest of a forming peloton.

As it is often easier to experimentally measure relative rather than absolute changes in densities and velocities, we here report the evolution of the motor density/roadblock density/motor velocity (*ρ*_m_(*x*)*/ρ*_rb_(*x*)*/v*_m_(*x*)) relative to its final value once pelotons are formed (*ρ*_m_*/ρ*_rb_*/v*_m_) (for details see The Supplementary Material)

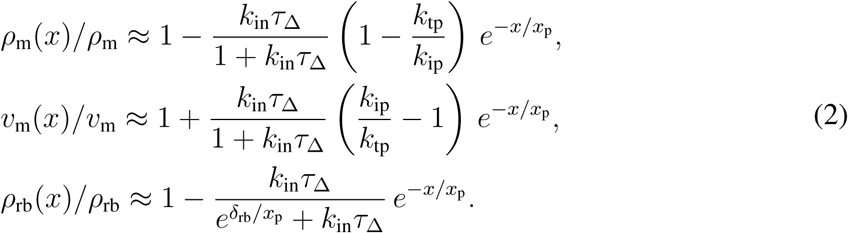

From Equation 2 we see that all measures approach their final value with an exponential decay over the region where pelotons form. We see that relative changes along the track grow in magnitude with the initiation rate, but that the effect saturates around *k*_in_ *∼* 1*/τ*_Δ_ for motor-density and velocity changes, while the roadblock density saturates later, around *k*_in_ *∼* e^*δ*_rb_/*x*_p_^ /*τ*_Δ_. The relative increase of the velocity and the densities saturate because when the initiation rate is larger than the effective roadblock binding time *k*_in_ *>* 1*/τ*_Δ_ most initiating motors have no roadblock in front of them and the initial density and velocity become independent of the initiation rate. The evolution of the density and velocity further depends on the strength of interactions between motors and roadblocks (*k*_tp_*/k*_ip_) since the change in velocity (density) when a motor catches up with a peloton is larger when *k*_tp_*/k*_ip_ is smaller.

In Figure 3 B we illustrate how the relative change in motor density is affected by the motor initiation rate, comparing the full expressions derived in The Supplementary Material to simulations. For slow initiation (*k*_in_*τ*_Δ_ *<* 1, green arrow in Figure 3 B), a roadblock typically binds between every two initiating motors. For such initiation rates, pelotons are typically of size one, giving only a marginal motor density change along the track (top density profile of Figure 3 D). For faster initiation (*k*_in_*τ*_Δ_ *>* 1, red arrow Figure 3 B), multiple motors bind before a roadblock rebinds to the start of the track, pelotons are larger than one, and we have a substantial increase in motor density as we move away from the initiation site (lower density profile of Figure 3 D). In Figure 3 C we compare our prediction for the distance over which pelotons form to estimates extracted by fitting an exponential relaxation distance to the density profiles generated by simulations. It is quite remarkable that our crude approximations capture the simulated data without any adjustable parameters.

## 3 Results

### 3.1 From pelotons to bursts

Transcriptional bursts have been observed in both eukaryotic and prokaryotic systems (*22, 42*), and are often ascribed to a promoter that can be turned on and off (*51*) (Figure 4 A). In the presence of roadblocks along the gene, our model shows that we should expect the same type of bursts even for promoters that are constantly turned on (Figure 4 B). To facilitate future experimental testing through the many known downstream effects of a bursty promoter (*51*), we here relate the bursts of motor activity in our model to those arising from the standard assumption of a promoter that turns on and off as described by a two-state model (*55*), (Figure 4 C). This should prove especially useful when characterizing the level of noise in mRNA production by using the Fano factor (the ratio between the variance and the mean in mRNA copy numbers), which has been widely used as a measure to classify transcriptional noise experimentally (*57, 59*). The fano factor for the two-state model is known (*51, 55*), such that a mapping of our model to the two-state model allows for a direct comparison with experiments.

**Figure 4:**
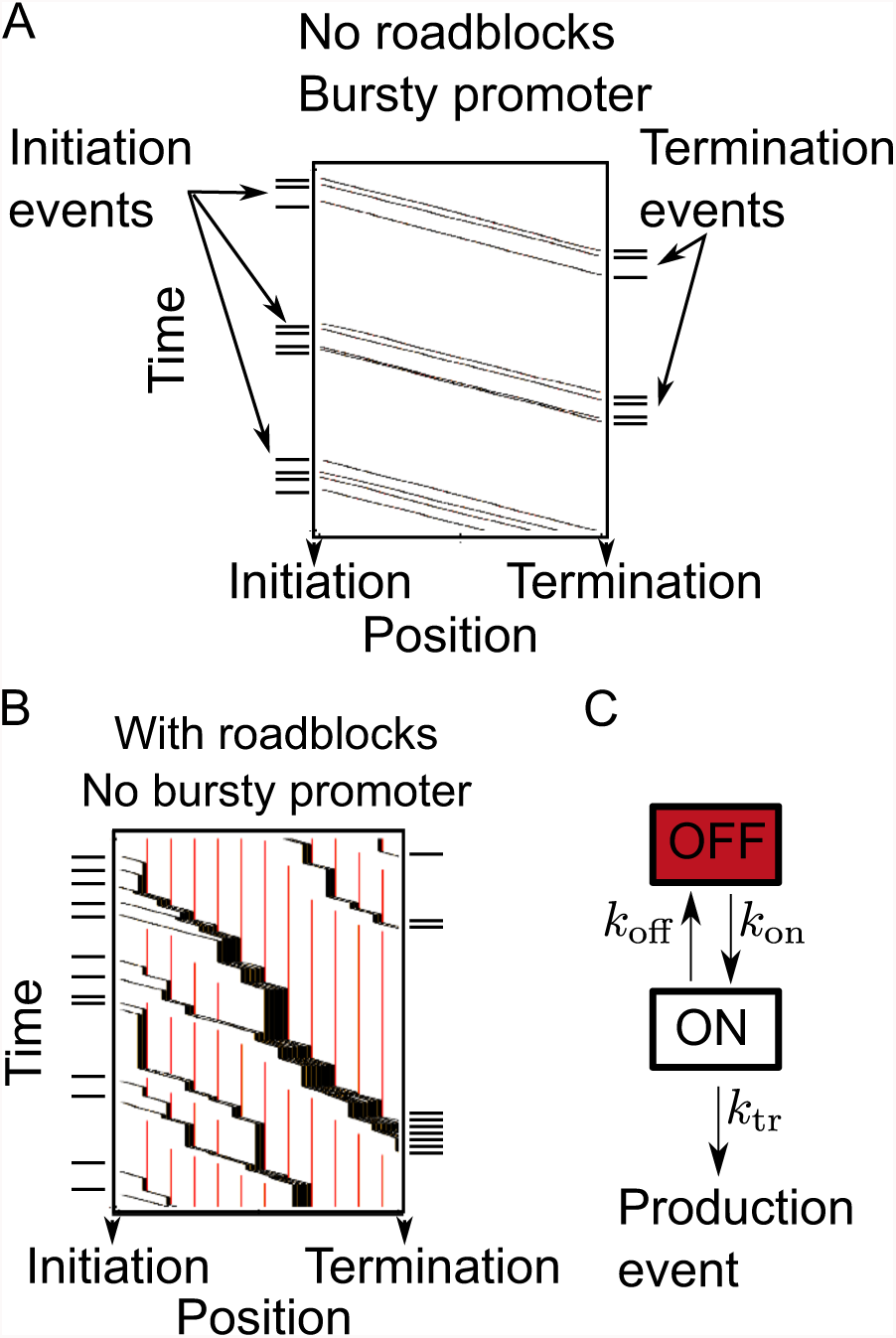
Burst generation from the promoter and during elongation. **(A)** A kymograph of a Monte Carlo simulation for transcription with a bursty promoter and no roadblocks. The time distribution of initiation events is similar to the time distribution at termination. **(B)** A kymograph for transcription without a bursty promoter, but with roadblocks along the gene (shown in red). Though the initiation events are exponentially distributed over time, the events at termination are more clustered, resulting in bursts of RNA production. **(C)** The phenomenological two-state model normally used to describe bursty transcription. In Equation 3 we report the parameters that would result from fitting the bursts generated by our model to the two-state model.

In the two-state model (Figure 4 C), the system switches between an on-state with production rate *k*_tr_, and an off-state where nothing is produced. The off-state switches to the on-state with rate *k*_on_, and back again with rate *k*_off_. Though we present the full form of how the effective burst parameters depend on microscopic parameters in The Supplementary Material, we here again limit ourselves to the physiologically relevant case where initiation rates are low enough that motors typically clear the initiation site between attempted initiation events,

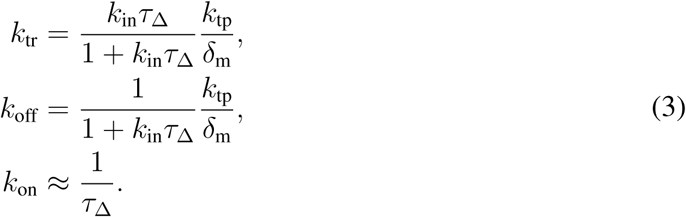

For significant peloton formation *k*_in_*τ*_Δ_ *>* 1 (Equation 1), the production rate *k*_tr_ becomes insensitive to the initiation rate, and is simply set by the rate at which motors pass termination. The off-rate *k*_off_ depends strongly on the initiation rate, and is given by the rate at which a typical peloton passes termination. The on-rate *k*_on_ is simply given by the inverse roadblock-rebinding time. It should also be noted that as long as the track extends further than the peloton forming distance *x*_p_ (which is the case for transcription, see below), the burst characteristics do not depend on the length of the track. The analytical relationships given in Equation 1-3 are the main results of this work. As these relationships dictate the precise dependence of a number of observables on microscopic parameters they should be well suited for falsification through comparison to future experiments (see Discussion). Next we show that the predictions are in line with the results from a number of recent studies, using as input parameters values from the literature (see Table 1).

### 3.2 Transcription on highly induced genes

Now that we have a quantitative understanding of how the non-specific interactions between motors and roadblocks give rise to peloton formation, we consider transcription on inducible genes in eukaryotes. These considerations are complicated by that nucleosome assembly and disassembly are not single step processes, as a tetramer and two dimers come together to make the full histone octamer contained in the nucleosome. *In vitro* studies have shown that a single polymerase only removes the histone dimer (*2, 3, 33*), while a second polymerase can dislodge the remaining hexamer (*37*). These *in vitro* results broadly agree with the *in vivo* observations that the density of histone dimers decreases strongly with transcription intensity genome wide, while an increased exchange and depletion of all core histones is only observed on highly transcribed genes (*10, 18, 29, 36, 40, 60, 69*).

On highly transcribed genes, we expext polymerases to form pelotons, and are thus expected to cooperate in dislodging the full nucleosome (*37*). Therefore, we assume that the roadblocks consist of histone hexamers that are dislodged by a passing polymerase. Further, we only consider genes where initiation is both active and non-paused, excluding situations where transcription is stalled by promoter-proximally paused polymerases (*12*). As Equation 1 shows that pelotons form when initiation is high (*k*_in_*τ*_Δ_ *>* 1), we compare our model to experiments tracking highly expressed genes (see Table 1).

We compare our analytical predictions of our heuristic approach to simulations (Figure 5). In our simulations we assume that the motors are only impeded at the nucleosome dyad, since this forms the largest obstacle for RNA polymerase II translocation (*26*). We consider the initiation rates *k*_in_ = 0.6 pol/min and *k*_in_ = 3.0 pol/min, where both rates correspond to highly induced genes, and the highest rate is chosen to match the maximal estimate of initiation rates on yeast genes (*56*). It is known that histones rebind on a sub-minute time scale (*63*), while it takes about a minute to clear space for a roadblock (see Table 1). Consequently, the nucleosome shadow is dominated by the roadblock size, and we assume Δ *≈ δ*_rb_ for simplicity in the analyticaltheory.

**Figure 5:**
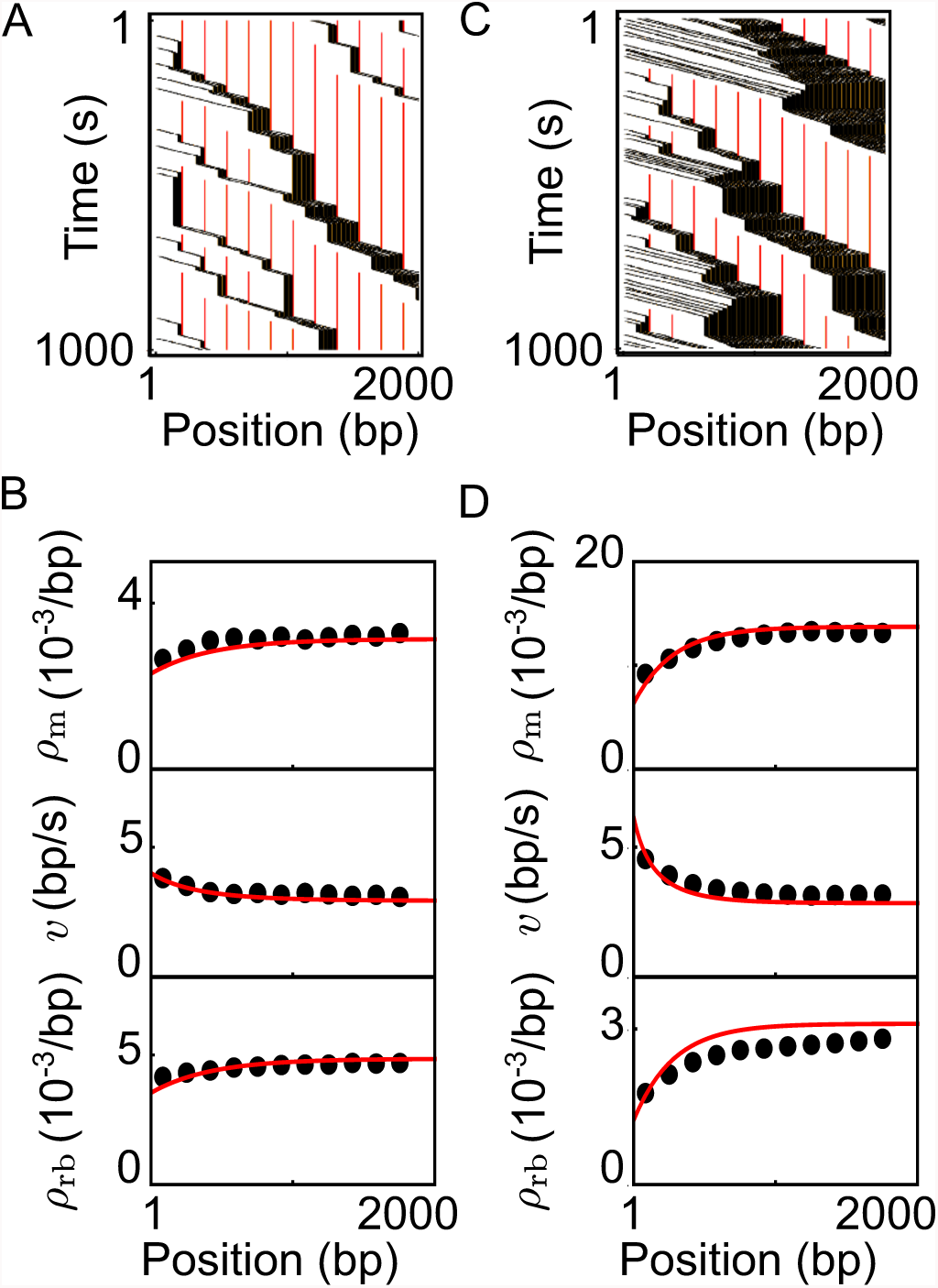
Bursts and density evolution for eukaryotic transcription on inducible genes. The parameter values used are shown in Table 1. **(A)** Kymograph for relatively moderate initiation rates. A polymerase (shown in black) evicts a nucleosome when it passes its center (the dyads, indicated by red lines). As the polymerases enter the gene, pelotons form over a distance of a few hundred base pairs. **(B)** The polymerase density, polymerase velocity, and nucleosome density corresponding to the kymograph in (A). Simulations were averaged over the size of a nucleosome and are shown as black dots, while our analytical predictions as red lines. **(C)** Kymograph for relatively high initiation rates, resulting in larger pelotons as compared to (A). **(D)** The polymerase density, polymerase velocity, and nucleosome density corresponding to the kymograph in (B). Comparing (B) and (D), we see that larger pelotons give a visibly stronger density and velocity evolution.

With only a small set of known microscopic input parameters (Table 1), our theory quantitatively captures the dynamics without free parameters. As predicted by Equation 1, we see that pelotons grow over the first few hundred base pairs after initiation (Figure 5 A and C). The peloton growth in turn means that the density of polymerases and nucleosomes near the initiation site is lower than further into the gene, while the velocity decreases as we move away from initiation (Figure 5 B and D). After the pelotons have formed, the polymerases and nucleosome densities, as well as polymerase velocities, remain virtually constant throughout the bulk of the gene. In Table 2 we give an overview of the predicted values of several observables, including burst parameters.

**Table 2:** Calculated observables for the physiological parameters in Table 1.

| **Initiation rate** | **0.6 pol/min** | **3.0 pol/min** |
| --- | --- | --- |
| $v_m(0)$: RNAP velocity at initiation | 5.5 bp/s | 8.1 bp/s |
| $v_m$: RNAP velocity once pelotons have formed | 3 bp/s | 3 bp/s |
| $J$: Transcriptional output | 0.6 mRNA/min | 2.3 mRNA/min |
| $\rho_m(0)$: RNAP density at initiation | 0.002 pol/bp | 0.006 pol/bp |
| $\rho_m$: RNAP density once pelotons formed | 0.003 pol/bp | 0.013 pol/bp |
| $n_p$: Peloton size | 1.6 pol | 3.8 pol |
| $x_p$: Distance over which pelotons form | 420 bp | 280 bp |
| $k_{on}$: Apparent on rate in two-state model | 1.1/min | 1.1/min |
| $k_{tr}$: Apparent production rate in two-state model | 1.8/min | 3.8/min |
| $k_{off}$: Apparent off rate in two-state model | 3.3/min | 1.4/min |

## 4 Discussion

With the aim of describing transcription in the crowded environment of the cell, we have introduced a general model that captures a large class of systems where molecular motors interact with dynamic roadblocks (Figure 1 B). Assuming that roadblocks have a finite size and/or rebinding time, and that motors slow down when evicting nucleosomes, we show that a physical mechanism reminiscent of drafting in racing sports gives rise to a strong kinetic attraction between motors. This attraction induces the motors to spontaneously reorganize into pelotons, and motors arrive to the terminus in bursts. Our analysis shows that one should always expect bursts in the presence of roadblocks if the motor initiation rate exceeds the inverse effective roadblock-rebinding time (*τ*_Δ_ in Equation 1), independently of if the promoter itself is bursty.

### 4.1 Peloton formation has been observed *in vivo*

Already 40 years ago there was evidence from Miller spreads suggesting that polymerases cluster on heavily transcribed genes (*1,25,39,50*). Recently, direct real-time evidence of polymerase ‘convoys’ on HIV-1 and POLR2A genes in HeLa cells has been produced (*67*). In accordance with our predictions, the typical distance between polymerases within convoys is too small for a nucleosome to bind, the distances between convoys are geometrically distributed, and a typical convoy includes several polymerases (see Table 2). Though our model can not rule out alternative explanations relying on specific polymerase-nucleosome interactions, the agreement between our predictions and the experimental observations without any adjustable fit parameters suggests that peloton formation through non-specific interactions with nucleosomes should be seriously considered, and could drive the formation of the observed polymerase convoys. This hypothesis can readily be tested by correlating the average peloton size (Equation 1) with changes in induction levels.

### 4.2 Predicted density profiles agree with observations in yeast

Our model also gives parsimonious explanations for several recent *in vivo* experimental observations pertaining to density profiles of polymerases and nucleosomes along inducible genes. In both yeast and human cells, highly transcribed genes without promoter-proximally paused polymerases show low nucleosome and polymerase densities for the first few hundred base pairs after initiation (*10, 13*). This distance agrees with our predicted distance over which are pelotons are formed, *x*_p_ (Table 2). Though there are many specific interactions that could give rise to nucleosome and polymerase depletion (*31*), the fact that this is a general property of heavily transcribed genes (*10, 16*) suggests a non-specific mechanism. Indeed, our model accurately predicts the occurrence and extent of such depletion without evoking any specific interactions (see Equation 1 and 2, as well as Figure 5 and Table 2). Our model correctly predicts a pausing-index (relative polymerase density within the promoter-proximal region compared to the bulk) that is below one for highly transcribed genes (*13*), and can be further tested by correlating changes in the pausing-index with for example histone modifications (*4*) that modify the transcription rate through nucleosomes.

The predicted increase of the polymerase density along the gene coincides with a decrease in the elongation rate (see Figure 5 B and D). Global run-on sequencing (GRO-seq) (*13*) experiments on active genes, on the contrary, have shown that the elongation rate *increases* over the first 15 kbp (*14*). The velocity increase in these GRO-seq experiments was likely caused by the gradual maturation of the transcription machinery (*31*) with mechanisms such as interactions with elongation factors not presently included in our model. Though it would be interesting to see how such mechanisms modulate the formation of pelotons, the observed increase of the elongation rate takes place over distances much longer than the few hundred base pairs over which pelotons form (see Table 2), and we do not expect our quantitative results to change due to these moderate velocity changes close to the promoter. Another source of velocity variation during transcription is that two polymerases are faster in evicting a nucleosome than one (*20*). We can see post hoc that such modifications would do little to change the model results, as we are interested in peloton forming systems where single polymerases forcing roadblocks are rare.

Another mechanism suggested to explain the observed density profiles is that transcription becomes termination-limited for high transcription rates (*10*). However, on a termination-limited gene, cued polymerases typically block each other's movement sterically (*48*) and leave no place for nucleosomes in between, which is inconsistent with experimental observations (*10*).

It is interesting to note that a nucleosome-free region at the start of genes (*10, 65, 74*) has been suggested to increase the accessibility of transcription factor binding sites close to the initiation site, thereby increasing the potential for transcriptional regulation (*65, 74*). Our model thus suggests that nucleosome depletion close to the initiation site could be a transcriptional epiphenomenon that has been coopted to allow for a greater regulatory response.

### 4.3 Burst characteristics agree with *in vivo* observations

Though bursts in RNA production has been observed in both prokaryotes (*7, 22*) and eukaryotes (*9, 57*), the origin is unclear, and usually modeled phenomenologically as arising from a promoter that turns on and off (*43, 58*). Though not developed with bursts in mind, our model predicts that transcriptions should be expected to be bursty as soon as the transcription initiation rate is comparable to the nucleosome binding rate, Equation 3, even if the promoter is constantly turned on. Several properties of the predicted bursts agree quantitatively with experimental observations. First, the pelotons are completed over a few hundred base pairs, which is shorter than the most genes. Therefore the predicted burst size is independent of gene length, agreeing with observations in yeast (*77*). Secondly, the predicted time between production events is on a sub-minute time scale (see Table 2), which falls within the experimentally observed range (*77*). Thirdly, our model predicts that when bursts are significant, only the apparent burst duration should be sensitive to transcription intensity, and that it decreases with increasing induction (see Equation 3). This behavior is broadly agreeing with the behavior reported for transcriptional bursting in *Escherichia coli* (*E. coli*) (*66*), where many other DNA binding proteins might act as the necessary roadblocks (*47*).

It should be noted that bursts generated during elongation does not rule out a bursty promoter. Instead, multi-scale bursting was recently reported (*67*) and could very well originate in a promoter turning on and off on long timescales, while pelotons form during elongation, giving rise to bursting on shorter timescales.

### 4.4 DNA supercoils as a source of bursts

Though there are many DNA binding proteins in *E. coli* (*47*), another interesting candidate for producing bursts is DNA supercoiling. Due to the helicity of DNA, transcribing polymerases are known to induce positive supercoils ahead and negative supercoils behind (*44*). Such supercoils slow down the polymerase (*7*), and will in the steady state extend some finite distance in front and behind. As negative supercoils spontaneously annihilate with positive supercoils, any DNA between two polymerases will have a lower net supercoiling density the closer together they are. With a lower supercoiling density ahead, a trailing polymerase will move faster than a leading polymerase, and all the conditions for peloton formation as described by our model (Figure 1 A) are fulfilled.

Our general mechanism of burst generation is connected to the mechanism suggested as a source of transcriptional bursting observed in bacteria, where a buildup of supercoils in torsional constrained plasmids was shown to suppress transcription until the supercoils were released (*7*). Importantly though, our model does not require the DNA to be torsional constrained as the supercoiling density around polymerases is set by the supercoils diffusivity (*72*) and a balance between supercoil creation and release.

### 4.5 Experimental testing and alternative models

Many mechanisms have been suggested for promoter induced bursting. As indicated in Figure 4, our model can be differentiated from such models by comparing the input and output dynamics. Our model could also be refuted by using existing techniques reporting on polymerase “convoys” (*67*) or transcriptional bursting (*22*) by manipulating or screening the limited set of effective parameters that controls the spatial and temporal evolution of the system (Equation 1-3). For example, the typical peloton size could be manipulated by changing the transcription initiation rate, or through histone modifications (*4*) that change the transcription rate through nucleosomes or the nucleosome rebinding time.

If the initiation dynamics cannot explain the bursts of RNA production, the elongation phase is likely the source of the bursts. To our knowledge, there are only two previous theoretical studies suggesting that bursts are created during elongation (*19, 73*). In both cases, intrinsic polymerase pausing through backtracking (*64*) was suggested as the source. However, back-tracking is unlikely to produce bursts, as it does not induce an effective attraction between polymerases, but rather an effective repulsion: interaction with a trailing polymerases is known to help terminate backtracks of a leading polymerase, and so speeds it up; interaction with a leading polymerase increases the chance of pausing in a trailing polymerase (*30, 37*), and so slows it down. Polymerases thus kinetically repulse each other, and jams induced by backtracks are unstable. Instead, we have shown that the interaction with roadblocks induces a persistent effective attraction between polymerases, resulting in a fast buildup of stable pelotons as polymerases move through the gene to terminate in bursts.

### 4.6 Conclusion and outlook

Our model points to a single source for a wide range of observed phenomena, from burst characteristics to the spatial organization of polymerases and nucleosomes. Surprisingly, the model agrees quantitatively with multiple experimental observations without adjustable parameters. Though further experiments are needed to determine the degree to which the observed phenomena can be explained through the non-specific polymerase and nucleosome interactions as we suggest, this work has the potential to reshape our understanding of how transcribing polymerases and nucleosomes organize spatially and temporally in physiologically crowded environments. Only by first understanding this organization, and how it can be modulated to effect things like polymerase cooperation, will it be possible to fully understand the action of transcription factors and other important cellular responses acting the elongation phase of transcription.

## Acknowledgements

We thank Joachim Griesenbeck for insightful discussions, Misha Klein, Orkide Ordu, Behrouz Eslami, John van Noort, and Stephan Grill for comments on the manuscript, and Ruben van Drongelen for programming support. MD and AAvdB acknowledge financial support from a TU Delft startup grant to MD. AAvdB further acknowledges financial support by the Netherlands Organization for Scientific Research (NWO/OCW), as part of the Frontiers of Nanoscience program.

## Author contributions

MD and AAvdB designed research, performed the research, and wrote the paper. AAvdB performed the Monte Carlo simulations and analyzed the simulation data.

## References

1. Benjamin Albert, Isabelle Léger-Silvestre, Christophe Normand, Martin K Ostermaier, Jorge Pérez-Fernández, Kostya I Panov, Joost C B M Zomerdijk, Patrick Schultz, and Olivier Gadal. RNA polymerase I-specific subunits promote polymerase clustering to enhance the rRNA gene transcription cycle. The Journal of cell biology, 192(2):277–93, 2011.

2. Dimitar Angelov, Vladimir a Bondarenko, Sébastien Almagro, Hervé Menoni, Fabien Mongélard, Fabienne Hans, Flore Mietton, Vasily M Studitsky, Ali Hamiche, Stefan Dimitrov, and Philippe Bouvet. Nucleolin is a histone chaperone with FACT-like activity and assists remodeling of nucleosomes. The EMBO journal, 25(8):1669–1679, 2006.

3. Rimma Belotserkovskaya, Sangtaek Oh, Vladimir a Bondarenko, George Orphanides, Vasily M Studitsky, and Danny Reinberg. FACT facilitates transcription-dependent nucleosome alteration. Science, 301(5636):1090–1093, 2003.

4. Lacramioara Bintu, Toyotaka Ishibashi, Manchuta Dangkulwanich, Yueh-Yi Wu, Lucyna Lubkowska, Mikhail Kashlev, and Carlos Bustamante. Nucleosomal elements that control the topography of the barrier to transcription. Cell, 151(4):738–49, nov 2012.

5. R A Blythe and M R Evans. Nonequilibrium steady states of matrix-product form: a solver's guide. Journal of Physics A: Mathematical and Theoretical, 40(46):R333–R441, nov 2007.

6. Kristin Brogaard, Liqun Xi, Ji-Ping Wang, and Jonathan Widom. A map of nucleosome positions in yeast at base-pair resolution. Nature, 486(7404):496–501, 2012.

7. Shasha Chong, Chongyi Chen, Hao Ge, and X. Sunney Xie. Mechanism of Transcriptional Bursting in Bacteria. Cell, 158(2):314–326, July 2014.

8. Debashish Chowdhury, Andreas Schadschneider, and Katsuhiro Nishinari. Physics of transport and traffic phenomena in biology: From molecular motors and cells to organisms. Physics of Life Reviews, 2(4):318–352, 2005.

9. Jonathan R Chubb, Tatjana Trcek, Shailesh M Shenoy, and Robert H Singer. Transcriptional Pulsing of a Developmental Gene. Current Biology, 16:1018–1025, 2006.

10. Hope A Cole, Josefina Ocampo, James R Iben, Răzvan V Chereji, and David J Clark. Heavy transcription of yeast genes correlates with differential loss of histone H2B relative to H4 and queued RNA polymerases. Nucleic acids research, 42(20):12512–22, November 2014.

11. Geoffrey M Cooper. The Cell: A Molecular Approach. 2nd edition. Sinauer Associates, 2000.

12. Leighton J Core and John T Lis. Transcription regulation through promoter-proximal pausing of RNA polymerase II. Science (New York, N.Y.), 319(5871):1791–2, mar 2008.

13. Leighton J Core, JJ. Waterfall, and John T. Lis. Nascent RNA Sequencing Reveals Widespread Pausing and Divergent Initiation at Human Promoters. Science, 322(December):1845–1848, 2008.

14. Charles G. Danko, Nasun Hah, Xin Luo, André L. Martins, Leighton Core, John T. Lis, Adam Siepel, and W. Lee Kraus. Signaling Pathways Differentially Affect RNA Polymerase II Initiation, Pausing, and Elongation Rate in Cells. Molecular Cell, 50(2):212–222, 2013.

15. Xavier Darzacq, Yaron Shav-Tal, Valeria de Turris, Yehuda Brody, Shailesh M Shenoy, Robert D Phair, and Robert H Singer. In vivo dynamics of RNA polymerase II transcription. Nature structural & molecular biology, 14(9):796–806, sep 2007.

16. P. P. Dennis, M. Ehrenberg, D. Fange, and H. Bremer. Varying rate of RNA chain elongation during rrn transcription in Escherichia coli. Journal of Bacteriology, 191(11):3740–3746, 2009.

17. B. Derrida, M R Evans, V Hakim, and V Pasquier. Exact solution of a 1D asymmetric exclusion model using a matrix formulation. Journal of Physics A: Mathematical and General, 26(7):1493–1517, apr 1993.

18. M F Dion, T Kaplan, M Kim, S Buratowski, N Friedman, and O J Rando. Dynamics of Replication-Independent Histone Turnover in Budding Yeast. Science, 315(March):1405–1409, 2007.

19. Maciej Dobrzynski and Frank J Bruggeman. Elongation dynamics shape bursty transcription. PNAS, 106(8):2583–2588, 2009.

20. Vitaly Epshtein and Evgeny Nudler. Cooperation Between RNA Polymerase Molecules in Transcription Elongation. Science, 300(5620):801–805, May 2003.

21. Ilya J Finkelstein and Eric C Greene. Molecular traffic jams on DNA. Annual review of biophysics, 42:241–63, jan 2013.

22. Ido Golding, Johan Paulsson, Scott M Zawilski, and Edward C Cox. Real-time kinetics of gene activity in individual bacteria. Cell, 123(6):1025–36, December 2005.

23. D S Goodsell. The Machinery of Life. Springer Sciences & Business Media, 2009.

24. Sandra J Greive and Peter H Von Hippel. Thinking quantitatively about transcriptional regulation. Nature reviews. Molecular cell biology, 6(3):221–32, mar 2005.

25. Francis Harper and Francine Puvion-Dutilleul. Non-Nucleolar Transcription Complexes of Rat Liver as Revealed by Spreading Isolated Nuclei. J Cell Sci, 40:181–192, 1979.

26. Courtney Hodges, Lacramioara Bintu, Lucyna Lubkowska, Mikhail Kashlev, and Carlos Bustamante. Nucleosomal fluctuations govern the transcription dynamics of RNA polymerase II. Science (New York, N.Y.), 325(5940):626–8, jul 2009.

27. J Howard. Mechanics of Motor Proteins and the Cytoskeleton. Sinauer Associates, Sunderland, Massachusetts, 2001.

28. M A Hoyt, A A Hyman, and M Bähler. Motor proteins of the eukaryotic cytoskeleton. Proceedings of the National Academy of Sciences of the United States of America, 94(November):12747–12748, 1997.

29. Adil Jamai, Rachel Maria Imoberdorf, and Michel Strubin. Continuous Histone H2B and Transcription-Dependent Histone H3 Exchange in Yeast Cells outside of Replication. Molecular Cell, 25(3):345–355, 2007.

30. Jing Jin, Lu Bai, Daniel S Johnson, Robert M Fulbright, Maria L Kireeva, Mikhail Kashlev, and Michelle D Wang. Synergistic action of RNA polymerases in overcoming the nucleosomal barrier. Nature structural & molecular biology, 17(6):745–52, June 2010.

31. Iris Jonkers and John T Lis. Getting up to speed with transcription elongation by RNA polymerase II. Nature reviews. Molecular cell biology, 16(3):167–177, 2015.

32. Mads Kaern, Timothy C Elston, William J Blake, and James J Collins. Stochasticity in gene expression: from theories to phenotypes. Nature reviews. Genetics, 6(6):451–64, June 2005.

33. Maria L Kireeva, Wendy Walter, Vladimir Tchernajenko, Vladimir Bondarenko, Mikhail Kashlev, and Vasily M. Studitsky. Nucleosome remodeling induced by RNA polymerase II: Loss of the H2A/H2B dimer during transcription. Molecular Cell, 9(3):541–552, 2002.

34. Stefan Klumpp. Pausing and Backtracking in Transcription Under Dense Traffic Conditions. Journal of Statistical Physics, 142(6):1252–1267, January 2011.

35. Stefan Klumpp and Terence Hwa. Stochasticity and traffic jams in the transcription of ribosomal RNA: Intriguing role of termination and antitermination. Proceedings of the National Academy of Sciences of the United States of America, 105(47):18159–64, nov 2008.

36. Arnold Kristjuhan and Jesper Q Svejstrup. Evidence for distinct mechanisms facilitating transcript elongation through chromatin in vivo. The EMBO journal, 23(21):4243–4252, 2004.

37. Olga I Kulaeva, Fu-Kai Hsieh, and Vasily M Studitsky. RNA polymerase complexes co-operate to relieve the nucleosomal barrier and evict histones. Proceedings of the National Academy of Sciences, 107(25):11325–30, June 2010.

38. A Kunwar, A John, K Nishinari, Andreas Schadschneider, and Debashish Chowdhury. Collective traffic-like movement of ants on a trail: dynamical phases and phase transitions. Journal of the Physical Society of Japan, 73(11):2979–2985, 2004.

39. Charles D. Laird and W. Yean Chooi. Morphology of transcription units in Drosophila melanogaster. Chromosoma, 58(2):193–218, 1976.

40. Cheol-Koo Lee, Yoichiro Shibata, Bhargavi Rao, Brian D Strahl, and Jason D Lieb. Evidence for nucleosome depletion at active regulatory regions genome-wide. Nature genetics, 36(8):900–5, aug 2004.

41. William Lee, Desiree Tillo, Nicolas Bray, Randall H Morse, Ronald W Davis, Timothy R Hughes, and Corey Nislow. A high-resolution atlas of nucleosome occupancy in yeast. Nature genetics, 39(10):1235–44, oct 2007.

42. Tineke Lenstra, Joseph Rodriguez, Huimin Chen, and Daniel R Larson. Transcription Dynamics in Living Cells *. Annu. Rev. Biophys, 45:25–47, 2016.

43. Gene-wei Li, X Sunney Xie, and Thomas Hirschfeld. Central dogma at the single-molecule level in living cells. Nature, 475:308–315, 2011.

44. L F Liu and J C Wang. Supercoiling of the DNA template during transcription. Proceedings of the National Academy of Sciences, 84(October):7024–7027, 1987.

45. O J O Loan, M R Evans, and M E Cates. Jamming transition in a homogeneous one-dimensional system: The bus route model. Physical Review E, 58(2):1404–1418, 1998.

46. K Luger, a W Mäder, R K Richmond, D F Sargent, and T J Richmond. Crystal structure of the nucleosome core particle at 2.8 Åresolution. Nature, 389(6648):251–260, 1997.

47. Martijn S. Luijsterburg, Malcolm F. White, Roel van Driel, and Remus Th. Dame. The Major Architects of Chromatin: Architectural Proteins in Bacteria, Archaea and Eukaryotes. Critical Reviews in Biochemistry and Molecular Biology, 43(6):393–418, 2008.

48. C T MacDonald, J H Gibbs, and A C Pipkin. Kinetics of Biopolymerization on Nucleic Acid Templates. Biopolymers, 6:1–25, 1968.

49. Jacek Mazurkiewicz, J Felix Kepert, and Karsten Rippe. On the mechanism of nucleosome assembly by histone chaperone NAP1. The Journal of biological chemistry, 281(24):16462–72, June 2006.

50. Steven L Mcknight and Oscar L Miller. Post-replicative D. melanogaster Nonribosomal Embryos Transcription Units in. Cell, 17(July):551–563, 1979.

51. Brian Munsky, Gregor Neuert, and Alexander van Oudenaarden. Using gene expression noise to understand gene regulation. Science, 336(6078):183–7, April 2012.

52. K P N Murthy and G M Schutz. Aging in two- and three-particle annihilation processes. Physical Review E, 57(2):1388–1394, 1998.

53. Keir C Neuman, Elio A Abbondanzieri, Robert Landick, Jeff Gelles, and Steven M Block. Ubiquitous Transcriptional Pausing Is Independent of RNA Polymerase Backtracking. Cell, 115:437–447, 2003.

54. Andrea Parmeggiani, T. Franosch, and E. Frey. Totally asymmetric simple exclusion process with Langmuir kinetics. Physical Review E, 70(4 2):046101, 2004.

55. J Peccoud and Bernard Ycart. Markovian modelling of gene product synthesis. Theoretical Population Biology, 48:222–234, 1995.

56. Vicent Pelechano, Sebastian Chavez, and Jose E. Perez-Ortin. A Complete Set of Nascent Transcription Rates for Yeast Genes. PloS one, 5(11):e15442, 2010.

57. Arjun Raj, Charles S Peskin, Daniel Tranchina, Diana Y Vargas, and Sanjay Tyagi. Stochastic mRNA Synthesis in Mammalian Cells. PLoS biology, 4(10), 2006.

58. Arjun Raj and Alexander van Oudenaarden. Nature, nurture, or chance: stochastic gene expression and its consequences. Cell, 135(2):216–26, October 2008.

59. Jonathan M Raser and Erin K O Shea. Control of Stochasticity in Eukaryotic Gene Expression Jonathan. Science, 304(5678):1811–1814, 2006.

60. Anne Rufiange, Pierre-É tienne Jacques, Wajid Bhat, François Robert, and Amine Nourani. Genome-Wide Replication-Independent Histone H3 Exchange Occurs Predominantly at Promoters and Implicates H3 K56 Acetylation and Asf1. Molecular Cell, 27(3):393–405, 2007.

61. Mamata Sahoo, Jiajia Dong, and Stefan Klumpp. Dynamic blockage in an exclusion process. Journal of Physics A: Mathematical and Theoretical, 48(1):015007, 2015.

62. Andreas Schadschneider, Debashish Chowdhury, and Katsuhiro Nishinari. Stochastic Transport in Complex Systems. Elsevier B.V., 2011.

63. Marc a Schwabish and Kevin Struhl. Evidence for eviction and rapid deposition of histones upon transcriptional elongation by RNA polymerase II. Molecular and cellular biology, 24(23):10111–7, dec 2004.

64. Joshua W Shaevitz, Elio a Abbondanzieri, Robert Landick, and Steven M Block. Back-tracking by single RNA polymerase molecules observed at near-base-pair resolution. Nature, 426(6967):684–7, dec 2003.

65. Sushma Shivaswamy, Akshay Bhinge, Yongjun Zhao, Steven Jones, Martin Hirst, and Vishwanath R. Iyer. Dynamic remodeling of individual nucleosomes across a eukaryotic genome in response to transcriptional perturbation. PLoS Biology, 6(3):0618–0630, 2008.

66. Lok-Hang So, Anandamohan Ghosh, Chenghang Zong, Leonardo a Sepúlveda, Ronen Segev, and Ido Golding. General properties of transcriptional time series in Escherichia coli. Nature genetics, 43(6):554–60, June 2011.

67. Katjana Tantale, Florian Mueller, Alja Kozulic-Pirher, Annick Lesne, Jean-Marc Victor, Marie-Cécile Robert, Serena Capozi, Racha Chouaib, Volker Bäcker, Julio Mateos-Langerak, Xavier Darzacq, Christophe Zimmer, Eugenia Basyuk, and Edouard Bertrand. A single-molecule view of transcription reveals convoys of RNA polymerases and multi-scale bursting. Nature Communications, 7:12248, 2016.

68. Sheila S. Teves, Christopher M. Weber, and Steven Henikoff. Transcribing through the nucleosome. Trends in Biochemical Sciences, 39(12):577–586, 2014.

69. Christophe Thiriet and Jeffrey J. Hayes. Replication-independent core histone dynamics at transcriptionally active loci in vivo. Genes and Development, 19(6):677–682, 2005.

70. Hugh Trenchard. The peloton superorganism and protocooperative behavior. Applied Mathematics and Computation, 270:179–192, 2015.

71. Francesco Turci, Andrea Parmeggiani, Estelle Pitard, M Carmen Romano, and Luca Ciandrini. Transport on a lattice with dynamical defects. Phy, 012705:1–8, 2013.

72. M T J van Loenhout, M V de Grunt, and Cees Dekker. Dynamics of DNA Supercoils. Science, 338(6103):94–97, 2012.

73. Margaritis Voliotis, Netta Cohen, Carmen Molina-Paŕıs, and Tanniemola B Liverpool. Fluctuations, pauses, and backtracking in DNA transcription. Biophysical journal, 94(2):334–48, jan 2008.

74. Assaf Weiner, Amanda Hughes, Moran Yassour, Oliver J Rando, and Nir Friedman. High-Resolution Nucleosome Mapping Reveals Transcription-Dependent Promoter Packaging. Genome Research, pages 90–100, 2010.

75. A Worcel, S Han, and M L Wong. Assembly of newly replicated chromatin. Cell, 15(3):969–977, 1978.

76. G-C Yuan, Liu Y-J, M F Dion, M D Slack, L F Wu, A J Altshuler, and O J Rando. Genome-Scale Identification of Nucleosome Positions in S. cerevisiae. Science, 309(5734):626–630, 2005.

77. Daniel Zenklusen, Daniel R Larson, and Robert H Singer. Single-RNA counting reveals alternative modes of gene expression in yeast. Nature structural & molecular biology, 15(12):1263–71, December 2008.

